# Yeast Knowledge Graphs Database for Exploring *Saccharomyces cerevisiae* and *Schizosaccharomyces pombe*

**DOI:** 10.1101/2024.12.04.626523

**Authors:** Mani R Kumar, Karthick Raja Arulprakasam, An-Nikol Kutevska, Marek Mutwil, Guillaume Thibault

## Abstract

Biomedical literature contains an extensive wealth of information on gene and protein function across various biological processes and diseases. However, navigating this vast and often restricted-access data can be challenging, making it difficult to extract specific insights efficiently. In this study, we introduce a high-throughput pipeline that leverages OpenAI’s Generative Pre-Trained Transformer Model (GPT) to automate the extraction and analysis of gene function information. We applied this approach to 84,427 publications on *Saccharomyces cerevisiae* and 6,452 publications on *Schizosaccharomyces pombe*, identifying 3,432,749 relationships for budding yeast and 421,198 relationships for *S. pombe*. This resulted in a comprehensive, searchable online Knowledge Graph database, available at <u>yeast.connectome.tools</u> and <u>spombe.connectome.tools</u>, which offers users extensive access to various interactions and pathways. Our analysis underscores the power of integrating artificial intelligence with bioinformatics, as demonstrated through key insights into important nodes like Hsp104 and Atg8 proteins. This work not only facilitates efficient data extraction in yeast research but also presents a scalable model for similar studies in other biological systems.

**HIGHLIGHTS:**

- Generated Yeast Knowledge Graphs from full-text research articles.
- Analyzed over 90,000 publications for *Saccharomyces* and *Schizosaccharomyces* species.
- Extracted millions of relationships using GPT-based natural language processing.
- Yeast Knowledge Graphs accessible through interactive web platforms and APIs.
- Advanced tool enabling insights into gene networks and functional interactions.

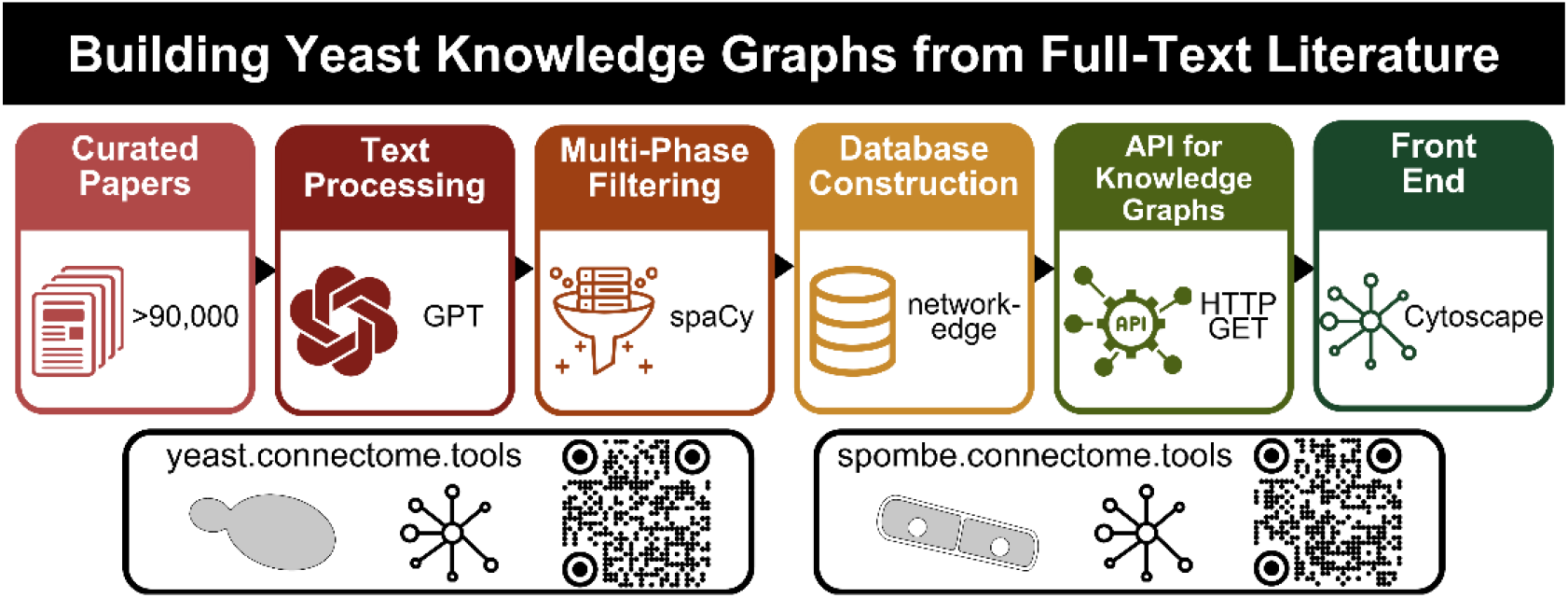

## INTRODUCTION

Research on yeast, particularly *Saccharomyces cerevisiae* and its fission counterpart, *Schizosaccharomyces pombe*, has expanded significantly in recent years. A search on PubMed reveals the scope of this growth: the number of publications on *S. cerevisiae* rose from 53,846 articles in the 2000s to 152,433 articles by 2023, while publications on *S. pombe* increased from 4,807 in the 2000s to 14,684 in 2023. This surge reflects the critical role of these model organisms in advancing our understanding of fundamental biological processes, from cell cycle regulation to stress responses and gene expression. However, the rapid pace of scientific publication makes it increasingly challenging for researchers to stay abreast of new discoveries, especially concerning gene and protein functions, interactions, and regulatory mechanisms.

Traditionally, researchers have relied on manually curated databases to organize and access experimentally validated “gold standard” data. Key resources, such as the *Saccharomyces* Genome Database (SGD) [1], BioGRID [2], and PomBase [3], have been instrumental in cataloging genetic and protein interactions, pathways, and functional annotations for *S. cerevisiae* and *S. pombe*. BioGRID, for instance, is a comprehensive repository for interaction datasets, facilitating the exploration of protein and genetic interactions across multiple organisms [2], like many databases, BioGRID and similar resources face inherent limitations, including delays in data curation due to the labor-intensive nature of manual annotation [4]. This poses a significant challenge, as researchers risk overlooking important insights within the constantly expanding corpus of yeast literature.

Manual curation, while thorough, is not scalable enough to keep pace with the influx of new information. The reliance on such methods results in a lag that limits researchers’ ability to understand the complex interactions underlying yeast biology in real time. This delay is particularly problematic in studying rapidly evolving areas, such as epigenetics, metabolic regulation, and cellular responses to environmental stressors. To bridge this gap, there is a pressing need for automated, high-throughput systems that can accelerate the integration of newly published data into accessible, structured repositories.

In response to this challenge, our study introduces an innovative approach that leverages the natural language processing capabilities of Generative Pre-trained Transformer (GPT) models. At the core of our method is a sophisticated text-mining pipeline powered by the GPT-3.5 turbo model from OpenAI. This pipeline systematically extracts and analyzes gene function information from a vast body of literature, covering abstracts and full-text articles on both *S. cerevisiae* and *S. pombe*. Through this high-throughput system, we have identified millions of relationships among genes, proteins, cellular compartments, environmental stresses, and other yeast-related entities, providing a comprehensive mapping of interactions within these model organisms.

Our Knowledge Graph databases—YeastKnowledgeGraph and FissionYeastKnowledgeGraph—are accessible online at yeast.connectome.tools and spombe.connectome.tools. These platforms offer an interactive interface that allows researchers to explore diverse interactions and pathways, while also providing an Application Programming Interface (API) for programmatic access. This resource not only democratizes access to yeast interaction data but also represents a scalable model for leveraging AI in bioinformatics, enabling faster and more efficient integration of scientific knowledge.

## RESULTS

### Semantic analysis and construction of Yeast Knowledge Graphs

Using the OpenAI GPT-3.5 model, we constructed Knowledge Graphs for two widely studied yeast model organisms: *S. cerevisiae* and *S. pombe*. Our approach involved processing 84,427 publications for *S. cerevisiae*, including 40,440 full-text articles and 43,987 abstracts, resulting in the creation of the YeastKnowledgeGraph—a comprehensive resource detailing 3,432,749 relationships among diverse yeast biological entities. For *S. pombe*, we processed 6,452 publications, comprising 1,089 full-text articles and 5,363 abstracts, which led to the development of the FissionYeastKnowledgeGraph with 421,198 documented relationships. Together, these publications spanned 1,867 and 457 journals for *S. cerevisiae* and *S. pombe*, respectively, covering a broad range of research areas relevant to yeast biology (Supplementary Table S1). To visually represent the scope of our data sources, we compiled the top 40 most frequently cited journals in each field, illustrating the quantitative distribution of articles (Figures 1A, B and Supplementary Table S1). These platforms serve as flexible tools for exploring biological entities, offering search capabilities based on keywords, author names, and PubMed IDs, as well as an ‘entities’ catalog listing all recognized entities within each database.

**Figure 1.**
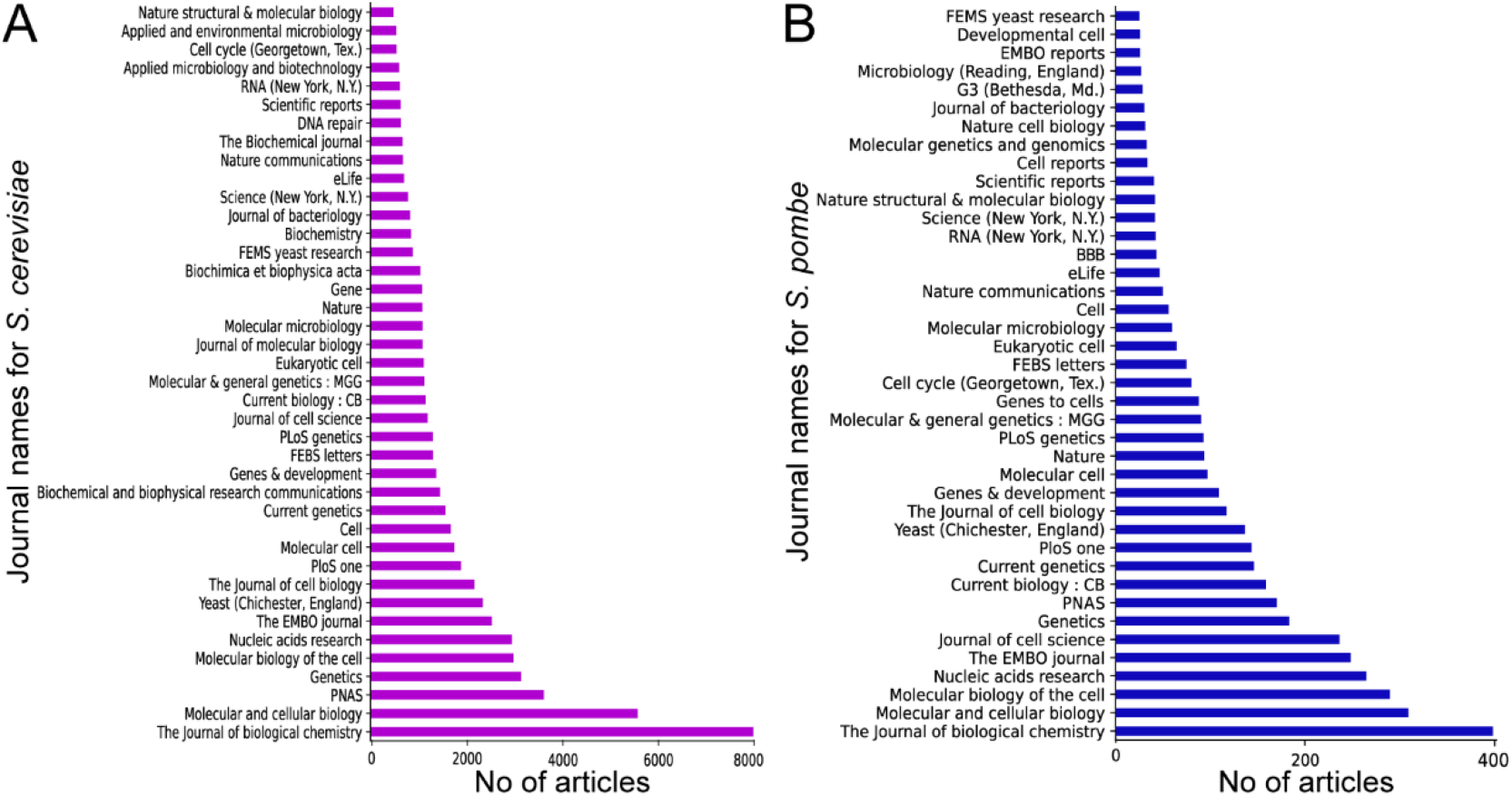
Distribution of Articles by Journal for *S. cerevisiae* and *S. pombe*. **A-B.** Quantitative distribution of articles from the top 40 journals publishing research on *S. cerevisiae* (A) and *S. pombe* **(B**).

Our analysis identified the 20 most frequently occurring entities, relationships, genes, and gene-to-gene interactions in both databases (Figures 2 and Supplementary Tables S2-S5). Within the YeastKnowledgeGraph, ‘*Saccharomyces cerevisiae*’ and CELLS emerged as prominent entities, whereas ‘*SWI6*’ and ‘*ATF1*’ were particularly notable in the FissionYeastKnowledgeGraph (Figure 2C-D). A common feature across both Knowledge Graphs was the predominance of ‘interacts with’ edges connecting entities (Figures 2A-B) and genes (Figures 2G-H). Key genes such as ‘*RAD51*’, RAD52, and HSP104 were extensively characterized in the YeastKnowledgeGraph due to their critical roles in yeast biology (Figure 2E). Similarly, ‘*SWI6*’, ‘*ATF1*’, and ‘*STY1*’ were highlighted as significant within the FissionYeastKnowledgeGraph (Figure 2F). This comparative analysis underscores the complexity and specificity of gene-to-gene interactions within these model organisms, providing a rich foundation for future research.

**Figure 2.**
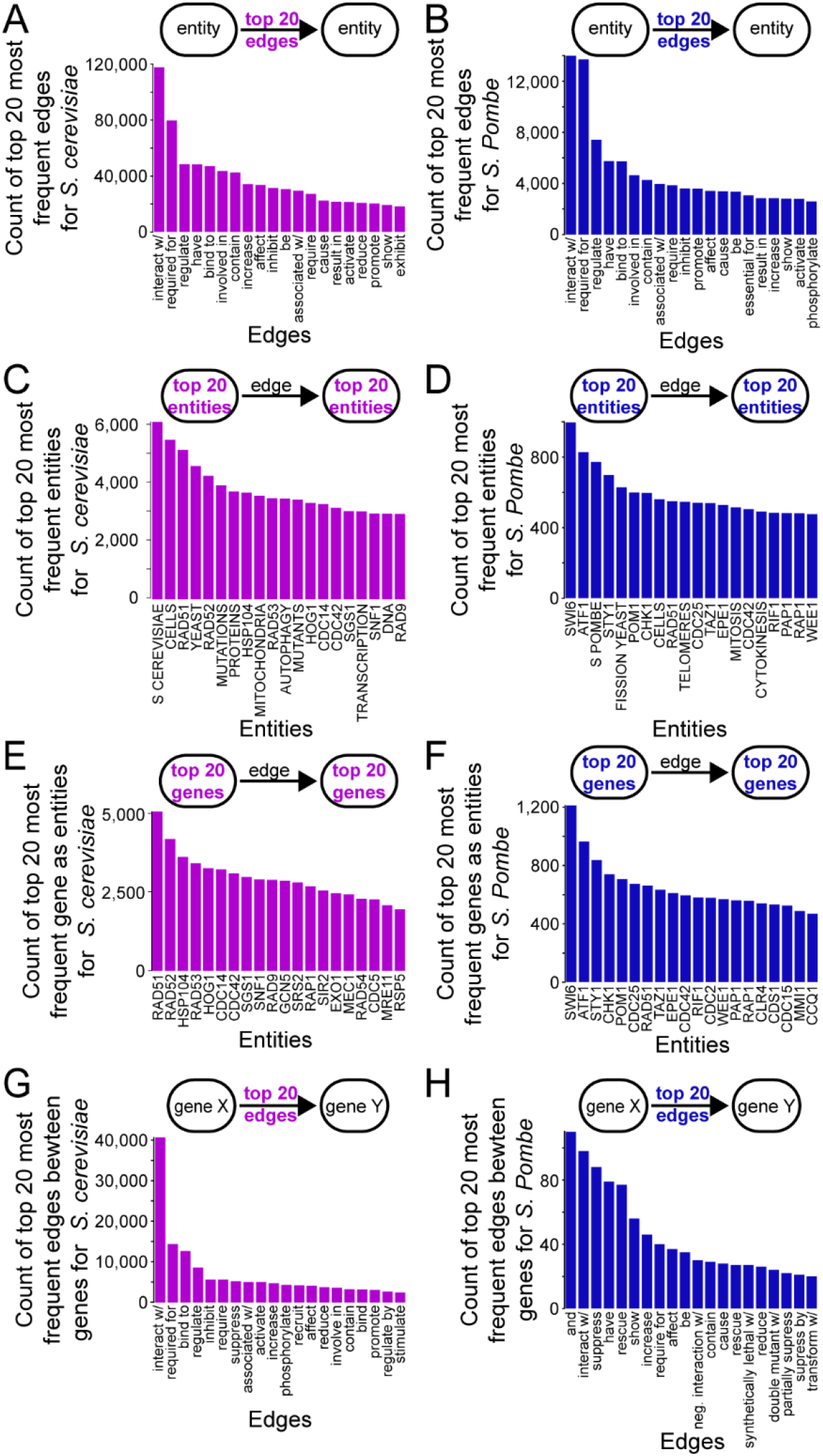
Network analysis highlights in *S. cerevisiae* and *S. pombe* Knowledge Graphs. **A-B.** Frequency distribution of the top 20 most frequent entities in the *S. cerevisiae* (A) and *S. pombe* (B) Knowledge Graphs. **C-D**. Frequency distribution of the top 20 frequently mentioned genes in the *S. cerevisiae* (C) and *S. pombe* (D) Knowledge Graphs. **E-F**. Frequency distribution of the top 20 most common interaction edges in the *S. cerevisiae* (E) and *S. pombe* (F) Knowledge Graphs. **G-H**. Frequency distribution of the top 20 most common interaction edges specifically between genes in the *S. cerevisiae* (G) and *S. pombe* (H) Knowledge Graphs.

### Interactive features and user engagement

The KnowledgeNetwork visualization offers users a dynamic, interactive experience with the Knowledge Graph, allowing them to engage directly with the network. By selecting nodes, users can activate tooltips that provide options for further exploration, such as examining neighboring nodes or accessing dedicated pages for specific features. The network is highly customizable—users can remove individual nodes or clusters, filter by specific relationship types (e.g., “binds to,” “links with,” “encodes for”), and adjust the layout for more focused analyses. To maintain clarity and readability, the network visualization is limited to 500 nodes.

For those requiring a more comprehensive dataset, a tab-delimited file of the entire network is available for download, enabling detailed offline analysis. Accompanying the KnowledgeNetwork visualization, a textual summary is provided, organized at the individual node level. This summary details each entity name, relationship type, and the corresponding PubMed IDs from which the data were extracted. This documentation aids users in validating relationships and addressing any potential inaccuracies inherent in GPT-based outputs. Additionally, users can view the network in a tabular format at the end of the results page for a straightforward, list-based perspective.

To extend the utility of the network for advanced research applications, web provide an API that enables programmatic access to the database. This API returns a JSON object containing relevant network and functional information, making it particularly valuable for researchers seeking to integrate the Knowledge Graph into computational workflows.

### Comparison of YeastKnowledgeGraph and BioGRID interactions

Our comparative analysis of BioGRID’s [4] protein-protein interaction (PPI) network with the YeastKnowledgeGraph dataset revealed that, out of 237,415 potential “interacts with” edges in the YeastKnowledgeGraph, 3,848 edges directly align with interactions cataloged in BioGRID (Figure 3A). YeastKnowledgeGraph also identified an additional 233,567 unique interaction edges not present in BioGRID, underscoring its enhanced detection capacity for diverse biological interactions. Furthermore, we observed 1,607 overlapping edges with different interaction types, such as “phosphorylates”, “inhibits”, and genetic interactions demonstrating the Knowledge Graph’s ability to categorize a variety of interaction types beyond traditional PPI data.

**Figure 3.**
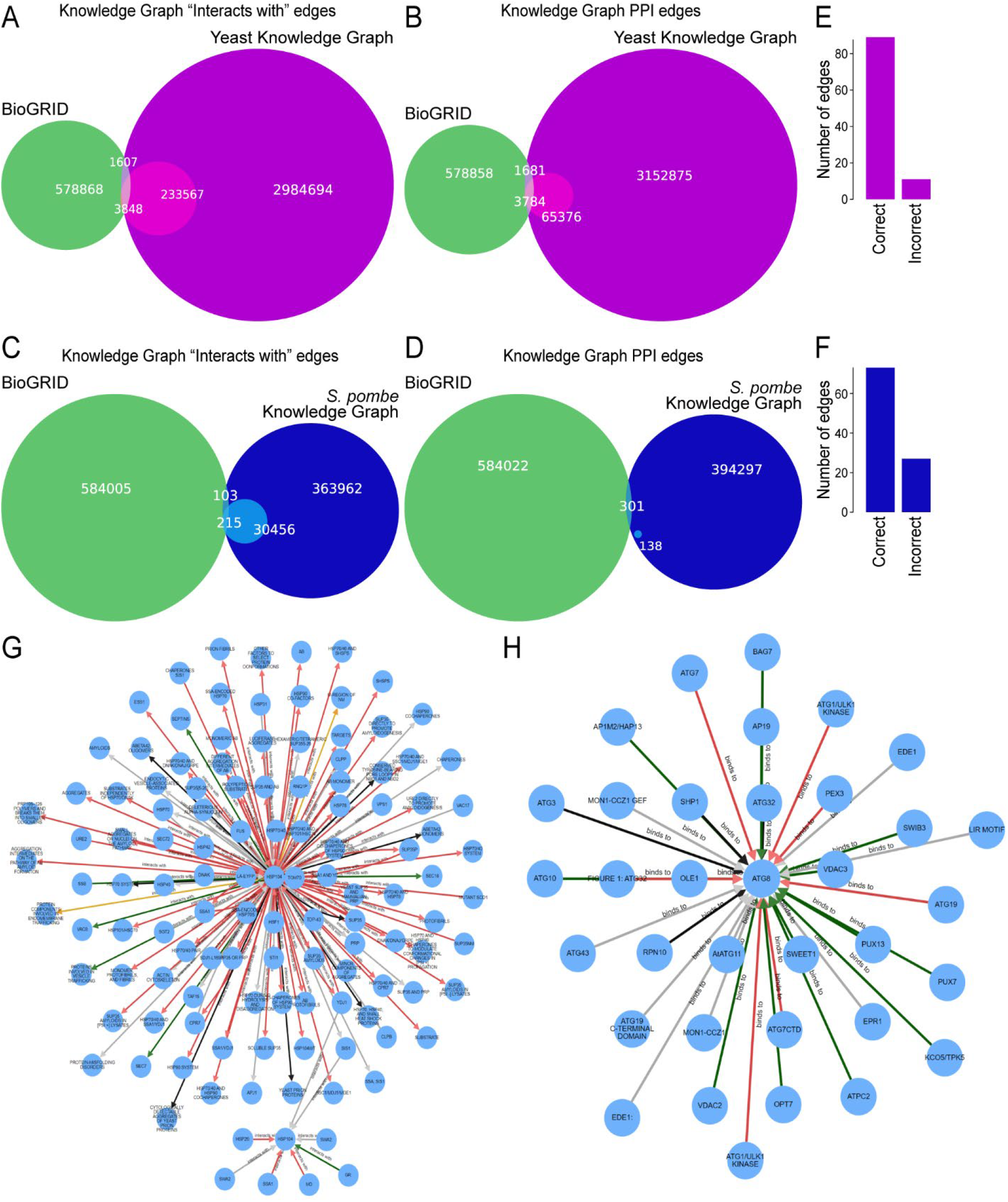
Comparative BioGRID Analysis and Key Gene Visualizations. **A-B.** Venn diagram illustrating the commonalities “interacting with” (A) and “PPI” (B) between edges in the *S. cerevisiae* Knowledge Graph and BioGRID’s interacting proteins dataset. **C-D**. Venn diagram illustrating the commonalities “interacting with” (C) and “PPI” (D) between edges in the *S. pombe* Knowledge Graph and BioGRID’s interacting proteins dataset. **E-F**. Bar chart indicating the accuracy assessment of randomly selected edges from the *S. cerevisiae* (E) and *S. pombe* (F) Knowledge Graphs. **G**. Knowledge network for the *S. cerevisiae* gene *HSP104*, with the “interacts with” relationship filter applied in Layout Options. This network highlights the various interactions *HSP104* has with other entities, providing insights into its role in cellular stress response. **H**. Knowledge network for the *S. cerevisiae* gene *ATG8*, with the “binds to” relationship filter applied in Layout Options. This network demonstrates the interactions of *ATG8*, especially in the context of autophagy-related binding interactions.

When focusing exclusively on PPI networks, YeastKnowledgeGraph showed an overlap of 3,784 PPI edges with BioGRID and identified 65,376 novel PPI edges not found in BioGRID (Figure 3B). These findings highlight YeastKnowledgeGraph’s capacity to comprehensively capture a broad spectrum of biological interactions, adding depth and diversity to the existing datasets in BioGRID.

### Comparison of FissionYeastKnowledgeGraph and BioGRID interactions

Expanding our analysis to the *S. pombe* Knowledge Graph, we found that 215 “interacts with” edges out of a total of 30,671 edges aligned with those in BioGRID (Figure 3C). The *S. pombe* Knowledge Graph further revealed 30,456 additional unique interaction edges, along with 138 novel PPI edges that were not cataloged in BioGRID (Figure 3D).

To assess the accuracy of the new edges identified within both the Yeast and *S. pombe* Knowledge Graphs, we conducted a manual review of a random sample of 100 edges. This evaluation demonstrated an 89% accuracy rate for edges within the YeastKnowledgeGraph and a 72% accuracy rate for edges in the *S. pombe* Knowledge Graph (Figure 3E-F and Supplementary Table S6). These findings confirm the Knowledge Graph’s reliability and underscore its potential as a valuable supplementary resource for the BioGRID database.

### Applications of YeastKnowledgeGraphs in Gene Regulatory Network Analysis

The YeastKnowledgeGraph serves as an invaluable tool for the yeast research community, providing a centralized repository of extensive data extracted from research abstracts and full-text articles. This resource enables researchers to explore gene regulatory networks, protein complexes, metabolic pathways, and stress responses. Here, we demonstrate the utility of the YeastKnowledgeGraph in dissecting the functional networks of specific proteins within *S. cerevisiae*.

#### Example 1 – YeastKnowledgeGraph Analysis of “*HSP104*”

Hsp104, a crucial protein disaggregase in *S. cerevisiae*, plays a significant role in thermotolerance by working with Hsp40 (Ydj1) and Hsp70 (Ssa1) to disassemble, resolubilize, and refold aggregated proteins under stress conditions [5]. The YeastKnowledgeGraph reveals the extensive network surrounding Hsp104. Querying “*HSP104*” generates a network map derived from 783 papers. Narrowing the focus to “interacts with” interactions through the “Layout Options” feature refines the network to 136 papers, with notable connections to Hsp40 (Ydj1) and Hsp70 (Ssa1), identified by terms such as “HSP70/40 PAIR,” “HSP SYSTEM,” “YDJ1,” and “SSA1.”

Interestingly, the Knowledge Graph also highlights Hsp104’s interactions with a variety of protein aggregate substrates, including “SUP35,” “LUCIFERASE AGGREGATES,” “ABETA42 MONOMERS,” and “PRION FIBRILS” (Figure 3G). This illustrates Hsp104’s broader involvement in protein homeostasis and stress response. While many of these interactions are well-documented, the YeastKnowledgeGraph provides a swift overview of the diverse conditions in which Sup35, among others, interacts with Hsp104. For example, specific interactions like “HSP104 interacts with HEXAMERIC/TETRAMERIC SUP355-26” [6] and “SOLUBLE SUP35” [7] are readily accessible through direct links to the publications, underscoring the utility of the Knowledge Graph in quickly retrieving detailed interaction information.

#### Example 2 – YeastKnowledgeGraph Analysis of “*ATG8*”

Atg8 (LC3), a ubiquitin-like protein essential for the formation of cytoplasm-to-vacuole transport vesicles and autophagosomes, plays a central role in the autophagy pathway [8–10]. The YeastKnowledgeGraph provides a comprehensive overview of Atg8’s interactions from a collection of 682 publications. Within this dataset, 41 papers describe Atg8 “binds to” interactions, shedding light on its dynamic role in the cellular environment (Figure 3H). Notably, Atg8 works in concert with Atf4 to facilitate the delivery of vesicles and autophagosomes to the vacuole via the microtubule cytoskeleton. The Knowledge Graph also underscores the roles of Atg3 and Atg7, both essential post-Atg4 for conjugating Atg8 with phosphatidylethanolamine (PE).

In contrast to conventional protein-protein interaction databases, the YeastKnowledgeGraph provides a more nuanced view of Atg8’s interactions, with direct links to the relevant literature. For instance, it details that Atg8 “binds to” the “C-TERMINAL FLEXIBLE TAIL OF ATG7” [11] and the “HR OF ATG3” [12]. Additionally, the Knowledge Graph reveals that Atg8 also “binds to” the “GROWING MEMBRANE” [13], contributing to a deeper understanding of its multifaceted role in cellular processes.

Furthermore, the Atg8 network captures its involvement in diverse biological processes beyond autophagy. For instance, it summarizes findings from 8 significant publications documenting Atg8’s role in “MACRONUCLEOPHAGY” [14], “HEMIFUSION OF LIPOSOMES” [8], “MEMBRANE TETHERING” [8], and “AGGREPHAGY” [15]. These insights demonstrate the Knowledge Graph’s ability to provide comprehensive and readily accessible information about key proteins in yeast biology.

## DISCUSSION

In this study, we introduce a novel application of advanced natural language processing, leveraging the OpenAI GPT-3.5 model to extract functional gene information from a vast collection of scientific literature. By processing over 90,879 abstracts and full-text articles, we developed the YeastKnowledgeGraph and Fission YeastKnowledgeGraph. These resources were built affordably (under USD1,000) and quickly (within two weeks), highlighting the efficiency of this approach in creating large-scale interaction databases. The resulting Knowledge Graphs encompass over 3.8 million relationships involving genes, proteins, cellular compartments, stress responses, and other yeast-related entities, making them valuable, interactive resources for the research community.

A key feature of the Yeast Knowledge Graph, the KnowledgeNetwork, provides an intuitive visual interface that allows researchers to filter specific relationships, focus on particular nodes, and personalize their exploration. This interactivity fosters a more targeted approach to data analysis, enabling researchers to streamline their investigation of complex biological networks. Our Knowledge Graphs demonstrate superior coverage and utility compared to established public repositories such as BioGRID[4], revealing numerous novel interactions and more granular interaction types (e.g., “phosphorylates,” “inhibits”). This added granularity underscores our Knowledge Graphs’ ability to capture a more comprehensive range of biological relationships, effectively complementing existing biological databases and filling critical knowledge gaps.

Despite the promising capabilities of our approach, some limitations remain. Access to literature, especially articles behind paywalls, restricts the scope of our data extraction and may lead to delays in incorporating the latest research findings. Furthermore, while automated techniques like GPT-3.5 offer scalability, they come with inherent trade-offs in precision when compared to manual curation. This is a known challenge in high-throughput text-mining pipelines. We are committed to refining our methodology and exploring enhanced access to publications to improve the accuracy and timeliness of our data.

In summary, our creation of the YeastKnowledgeGraph and FissionYeastKnowledgeGraph demonstrates the potential of generative AI models in advancing biological research. These Knowledge Graphs, accessible through an interactive, web-based platform, provide a valuable resource for understanding gene interactions and biological pathways in yeast. Our work lays the foundation for further advancements in the study of complex biological systems, with potential applications extending to other model organisms and beyond.

## LIMITATION OF THE STUDY

This study acknowledges several limitations that impact the scope and accuracy of our Knowledge Graphs. First, our reliance on publicly available literature and articles accessible through institutional subscriptions may result in a delay in incorporating the latest research findings, particularly those behind paywalls. This dependence limits the immediacy with which emerging data can be integrated into our resource, potentially affecting its relevance in rapidly evolving research areas.

Additionally, using a pre-trained model like GPT-3.5 in processing specialized scientific literature introduces certain challenges. Although GPT-3.5 offers significant capabilities in language processing, it occasionally struggles with producing methodologically structured and coherent results when applied to complex scientific texts. This can impact the clarity and precision of the resulting knowledge network. Another limitation stems from the model’s occasional misidentification of entities and relationships, which can affect the reliability of the analysis, particularly in dense, intricate sections of text.

To address these challenges, we are considering fine-tuning the language model with a targeted corpus of scientific research articles. Fine-tuning could improve the model’s comprehension of domain-specific terminology and its handling of complex biological relationships, resulting in a more accurate and coherent knowledge representation.

We are also evaluating the adoption of the more advanced GPT-4 model, which has demonstrated improved accuracy and proficiency in handling complex tasks due to an additional six months of training with human and automated feedback. For example, GPT-4 has shown enhanced accuracy rates in various predictive tasks, which could benefit our project. However, the use of GPT-4 comes at a significantly higher cost—approximately 20 times that of GPT-3.5. Therefore, our choice between these models will require a careful cost-benefit analysis, balancing budgetary constraints with our need for higher accuracy.

In future iterations, we aim to refine our methods further, optimize access to cutting-edge publications, and explore advanced model configurations to enhance both the quality and utility of the Knowledge Graphs. By addressing these limitations, we strive to ensure that our resource remains a robust, accurate, and timely tool for the yeast research community.

## METHODS

### Retrieval and pre-processing of literature

To build our comprehensive dataset, we obtained an extensive list of yeast genes and their aliases from the YeastMine database and UniProt. Using these lists, we created specialized search queries to locate relevant research articles in PubMed. For *Saccharomyces cerevisiae*, our search query took the form “(Saccharomyces cerevisiae[Title/Abstract] AND {gene}[tw])”, with a similarly structured query for *Schizosaccharomyces pombe*. These searches were automated using the Bio.Entrez package (v1.81), enabling efficient retrieval of articles containing information about gene functions in these organisms. Where available, we also retrieved full-text articles through the Elsevier API, supplementing our collection with abstracts for a more complete dataset.

### Processing of texts using GPT-3.5 Turbo

We utilized Python scripts to extract gene-specific information from the collected articles, employing OpenAI’s GPT-3.5-turbo model for text processing. The model’s API was configured with a temperature setting of zero to prioritize accuracy by minimizing randomness, thereby reducing the risk of misinformation. The model was tasked with identifying relevant entities, such as genes and proteins, and elucidating their relationships within the text. We designed a tailored prompt (Supplementary Table S7) to structure the output, ensuring that each line highlighted a pair of entities and their specific relationship, thereby optimizing the data for more detailed analysis.

### Multi-phased Filtering and Database Construction

Following the initial generation of relationship edges by the GPT-3.5-turbo model [16], a multi-phased filtering process was applied to validate and enhance the dataset. Using the spaCy [17] NLP library, we validated the generated edges to exclude any hallucinated edges—relationships that were inaccurately inferred and did not appear in the original text. Any edges with incorrect punctuation were marked as “bad edges” and re-submitted to the GPT model for correction. After this refinement process, edges were classified as either “good” for accurate entries or retained as “bad” if they continued to exhibit issues.

The finalized edges were subsequently used to build a network-edge database. We implemented the front-end application using the Python-Flask framework (v2.2.3), with Networkx (v3.1) handling graph analyses, and MongoDB (in conjunction with PyMongo) facilitating robust database management. To enhance the visualization experience, we incorporated Cytoscape.js (v3.23). This integrated approach ensured a reliable and well-organized dataset that could support interactive queries and efficient data visualization.

### API for yeast Knowledge Graphs

Both the Yeast and Fission Yeast Knowledge Graphs feature a robust Application Programming Interface (API) that allows remote search queries and data retrieval through HTTP GET requests. The API, built with the same technologies mentioned above, returns a JSON object containing relevant nodes, edges, and summarized texts in response to user queries. Users can access the API by appending /api/<search type>/<search query> to the base URL, where <search type> specifies the search category, and <search_query> represents the actual search term. This interface streamlines access to the Knowledge Graph data, facilitating deeper user engagement and enabling programmatic access to support a wide range of research applications.

## Data and Code Availability

The custom code to generate the biomaps is available at GitHub (https://github.com/mutwil/plant_connectome).

## Competing interests

The authors declare no competing financial interests.

## Funding

This work was supported by funds from the Singapore Ministry of Education Academic Research Fund Tier 1 (RG96/22 to G.T.) and Tier 3 (MOET32022-0002 to M.M.) as well as the Research Scholarship to K.R.A [predoctoral fellowship from Singapore Ministry of Education Academic Research Fund Tier 3 (MOE-MOET32020-0001)].

## Author contributions

Conceptualization: M.M. and G.T.; Methodology: M.M. and G.T.; Formal analysis: M.K.R., K.R.A., and A.N.K.; Investigation: M.R.K., K.R.A., and A.N.K.; Writing - original draft: M.R.K., G.T. and M.M.; Writing - review & editing: M.R.K., K.R.A., M.M., and G.T.; Supervision: M.M. and G.T.; Project administration: M.M. and G.T.; Funding acquisition: M.M. and G.T.

## Acknowledgements

We thank members of Mutwil and Thibault labs for critical reading of the manuscript.

## Notes

http://yeast.connectome.tools/

http://spombe.connectome.tools/

### Competing Interest Statement

The authors have declared no competing interest.

