## Supplementary Table S7 for "Yeast Knowledge Graphs Database for Exploring *Saccharomyces cerevisiae* and *Schizosaccharomyces pombe*"

**Supplementary Table S2.** An example of an abstract, prompts and outputs from GPT.

| | |
|---|---|
| Abstract | Gliotoxin (GT) is a secondary fungal metabolite with pleiotropic immunosuppressive properties that have been implicated in Aspergillus virulence. However, the mechanisms of GT cytotoxicity and its molecular targets in eukaryotic cells have not been fully characterized. We screened a haploid library of Saccharomyces cerevisiae single-gene deletion mutants (4,787 strains in EUROSCARF) to identify nonessential genes associated with GT increased resistance (GT-IR) and increased sensitivity (GT-IS). The susceptibility of the wild-type parental strain BY4741 to GT was initially assessed by broth microdilution methods using different media. GT-IR and GT-IS were defined as a fourfold increase and decrease, respectively, in MIC, and this was additionally confirmed by susceptibility testing on agar yeast extract-peptone-glucose plates. The specificity of GT-IR and GT-IS mutants exhibiting normal growth compared with the wild-type strain was further tested in studies of their susceptibility to conventional antifungal agents, cycloheximide, and H2O2. GT-IR was associated with the disruption of genes acting in general metabolism (OPI1, SNF1, IFA38), mitochondrial function (RTG2), DNA damage repair (RAD18), and vesicular transport (APL2) and genes of unknown function (YGL235W, YOR345C, YLR456W, YGL072C). The disruption of three genes encoding transsulfuration (CYS3), mitochondrial function (MEF2), and an unknown function (YKL037W) led to GT-IS. Specificity for GT-IR and GT-IS was observed in all mutants. Importantly, the majority (69%) of genes implicated in GT-IR (6/10) and GT-IS (2/3) have human homologs. We identified novel Saccharomyces genes specifically implicated in GT-IR or GT-IS. Because most of these genes are evolutionarily conserved, further characterization of their function could improve our understanding of GT cytotoxicity mechanisms in humans. |
| Prompt | messages= [<br>{"role": "system", "content": "You are data scientist tasked with identifying entities and relationships in scientific text. Please provide only one statement per line, and ensure that each line contains exactly two entities. If a relationship involves more than two entities, please break it down into multiple separate lines. "},<br>{"role": "user", "content": "CLas inhibits callose deposition in the sieve pores and the accumulation of reactive oxygen species to favor its cell-to-cell movement. CESA1 interacts with CESA4 and 7. HIS3 mutant inhibits proline production."},<br>{"role": "assistant", "content": "CLas: !inhibits! callose deposition\nCLas: !inhibits! accumulation of reactive oxygen species\ncallose deposition: !inhibits! cell-to-cell movement\nCESA1!interacts with!CESA4\nCESA1!interacts with!CESA7\nHIS3 mutant!inhibits!proline production"},<br>{"role":"user","content":"VERY SHORT, CONCISE SUMMARY CONTAINING ALL INFORMATION WITH TWO ACTORS PER LINE: "+text}<br>] |
| Output | GT-IR!associated with!OPI1<br>GT-IR!associated with!SNF1<br>GT-IR!associated with!IFA38<br>GT-IR!associated with!RTG2<br>GT-IR!associated with!RAD18<br>GT-IR!associated with!APL2<br>GT-IR!associated with!YGL235W<br>GT-IR!associated with!YOR345C<br>GT-IR!associated with!YLR456W<br>GT-IR!associated with!YGL072C<br>CYS3!disruption led to!GT-IS<br>MEF2!disruption led to!GT-IS<br>YKL037W!disruption led to!GT-IS<br>Specificity for GT-IR!observed in!all mutants<br>Specificity for GT-IS!observed in!all mutants |
